# Presence or absence of a prefrontal sulcus is linked to reasoning performance during child development

**DOI:** 10.1101/2022.03.02.482563

**Authors:** Ethan H. Willbrand, Willa I. Voorhies, Jewelia K. Yao, Kevin S. Weiner, Silvia A. Bunge

**Affiliations:** Department of Psychology, University of California Berkeley, Berkeley, CA, 94720 USA; Helen Wills Neuroscience Institute, University of California Berkeley, Berkeley, CA, 94720 USA; Princeton Neuroscience Institute, Princeton University, Princeton, NJ, 08540 USA

**Keywords:** neuroanatomy, neurodevelopment, brain imaging, sulcal pattern, cortical folding, prefrontal cortex, reasoning

## Abstract

The relationship between structural variability in late-developing association cortices like the lateral prefrontal cortex (LPFC) and the development of higher-order cognitive skills is not well understood. Recent findings show that the morphology of LPFC sulci predicts reasoning performance; this work led to the observation of substantial individual variability in the morphology of one of these sulci, the para-intermediate frontal sulcus (pimfs). Here, we sought to characterize this variability and assess its behavioral significance. To this end, we identified the pimfs in a developmental cohort of 72 participants, ages 6-18. When controlling for age, the presence or absence of the ventral component of the pimfs was associated with reasoning, as was the total surface area of pimfs. These findings show that multiple features of sulci can support the development of complex cognitive abilities and highlight the importance of considering individual differences in local morphology when exploring the neurodevelopmental basis of cognition.

## Introduction

A major interest in cognitive neuroscience is to understand how variability in brain structure relates to individual differences in complex cognition. Of the many cortical features to study, one of the most prominent is the patterning of the indentations, or sulci. While the majority of sulci are identifiable across individuals, some tertiary sulci that emerge late in gestation are not (Amiez et al., 2019; Chiavaras and Petrides, 2000; Lopez-Persem et al., 2019; Voorhies et al., 2021; Willbrand et al., 2021). For example, the variable presence of the paracingulate sulcus in the anterior cingulate cortex is related to performance on cognitive, motor, and affective tasks in young adults (Amiez et al., 2018; Buda et al., 2011; Fornito et al., 2006, 2004; Huster et al., 2011, 2009; Whittle et al., 2009), inhibitory control in children (Borst et al., 2014; Cachia et al., 2014), and in disorders such as schizophrenia (Le Provost et al., 2003; Yücel et al., 2003, 2002), obsessive-compulsive disorder (Shim et al., 2009), and frontotemporal dementia (Harper et al., 2022). Additional recent work also shows that i) sulci are missing in orbitofrontal cortex in individuals with schizophrenia or autism spectrum disorder (Nakamura et al., 2020) and ii) the absence of multiple sulci in the ventromedial prefrontal cortex (PFC) affects the functional organization of the default mode network (Lopez-Persem et al., 2019). However, it is presently unknown whether the presence or absence of tertiary sulci in lateral prefrontal cortex (LPFC) impacts cognition in child development.

For example, in a recent study (Voorhies et al., 2021), we demonstrated that the depth of specific LPFC tertiary sulci was related to abstract reasoning ability, or the ability to solve novel problems. We observed pronounced variability in the number of components, and overall prominence, of one of the three sulci identified by the model (para-intermediate frontal sulcus, pimfs). Here, we build on this observation by testing the targeted hypothesis that the presence or absence of specific components of the pimfs, and/or the prominence – quantified as the total sulcal surface area – of this sulcus was related to reasoning abilities. To do so, we investigated the relationship between the variability of the pimfs and relational reasoning scores in a sample of 72 children and adolescents aged 6-18.

## Results

Sulci were manually identified as component(s) of the pimfs in each hemisphere according to previous work (Fig 1A; Supplementary Fig. 1; Amiez and Petrides, 2007; Petrides, 2019; Voorhies et al., 2021). We confirmed our prior observation that the pimfs was highly variable (Fig. 1B): in a given hemisphere, there could be zero, (*left* = 2.78% of participants; *right* = 8.33%), one (*left*: 22.22%; *right*: 25%), or two components (*left*: 75%; *right*: 66.67%; *left: X^2^* = 60.333, df = 2, *p* < 0.001; *right: X^2^* = 39, df = 2, *p* < 0.001; no hemispheric asymmetry: *p* = 0.30). Based on published criteria (Petrides, 2013), we further defined pimfs components as either dorsal (pimfs-d) or ventral (pimfs-v) and assessed the prevalence of each component (Fig. 1B). Numerically, a single dorsal component was more frequent than a single ventral one, but statistically these profiles were equally frequent (*X^2^* ≥ .89,*p* > .30 for both hemispheres).

**Fig. 1.**
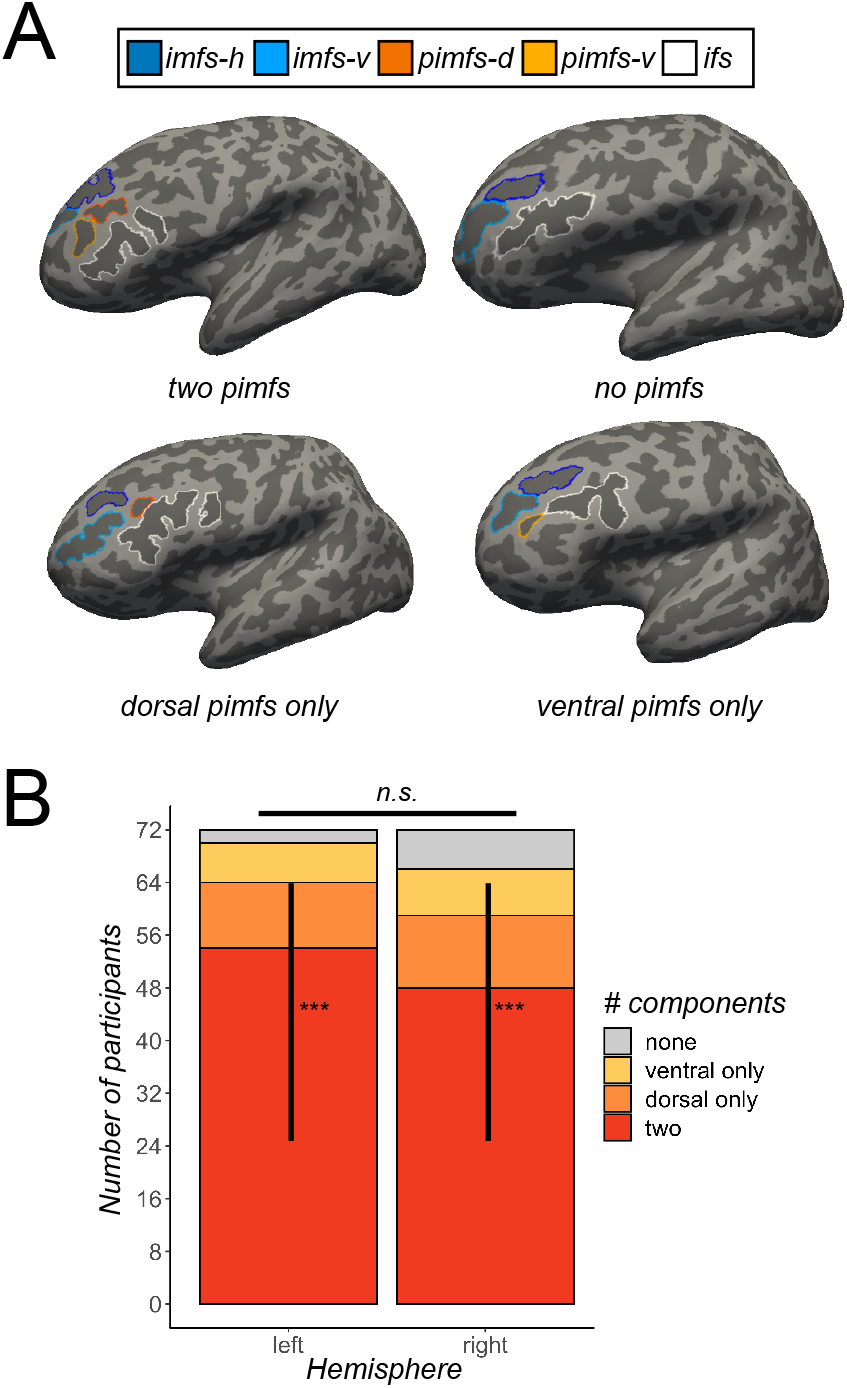
The para-intermediate middle frontal sulcus: A tertiary sulcus in lateral prefrontal cortex with extensive individual differences. (A) Four inflated left hemispheres (sulci: dark gray; gyri: light gray; Cortical surfaces are not to scale) depicting the four types of incidence of the para-intermediate middle frontal sulcus (pimfs): i) both components present, ii) neither present, iii) dorsal component present, iv) ventral component present. The prominent sulci bounding the pimfs are also shown: the horizontal (imfs-h) and ventral (imfs-v) intermediate middle frontal sulci and inferior frontal sulcus (ifs). These four sulci are colored according to the legend. (**B**) Stacked bar plot depicting the incidence of the pimfs components in both hemispheres across the sample (N=72 participants). The incidence of the pimfs is highly variable. In each hemisphere, it is more common for participants to have two components than a single component or no component (*** *p*s < .0001); the distribution of incidence does not differ between hemispheres (n.s., *p* = .3). When only one component was present in a hemisphere, it was equally likely to be a dorsal or ventral component (*p*s > .3).

An analysis of covariance (ANCOVA; including age as a covariate) testing the effect of pimfs incidence rates on reasoning found that the presence of two pimfs components in the left hemisphere was related to improved reasoning performance (F(1,67) = 4.18, *p* = 0.045, η2G = 0.059), but not the right hemisphere (F(1,67) = 2.63, *p* = 0.11, η2G = 0.038; Fig. 2A, left). An ANCOVA testing whether the incidence of a dorsal and/or ventral pimfs component, specifically, was linked to reasoning revealed that the presence of the left hemisphere pimfs-v was associated with higher reasoning scores, controlling for age (F(1,66) = 5.10, *p* = 0.027, η2G = 0.072; Supplementary Fig. 2). Follow-up analyses with additional behavioral measures revealed that the presence of the left pimfs-v was not related to processing speed or phonological working memory (digit span forwards and backward; all *p*s > 0.5), suggesting some degree of specificity in this brain-behavior relation.

**Fig. 2.**
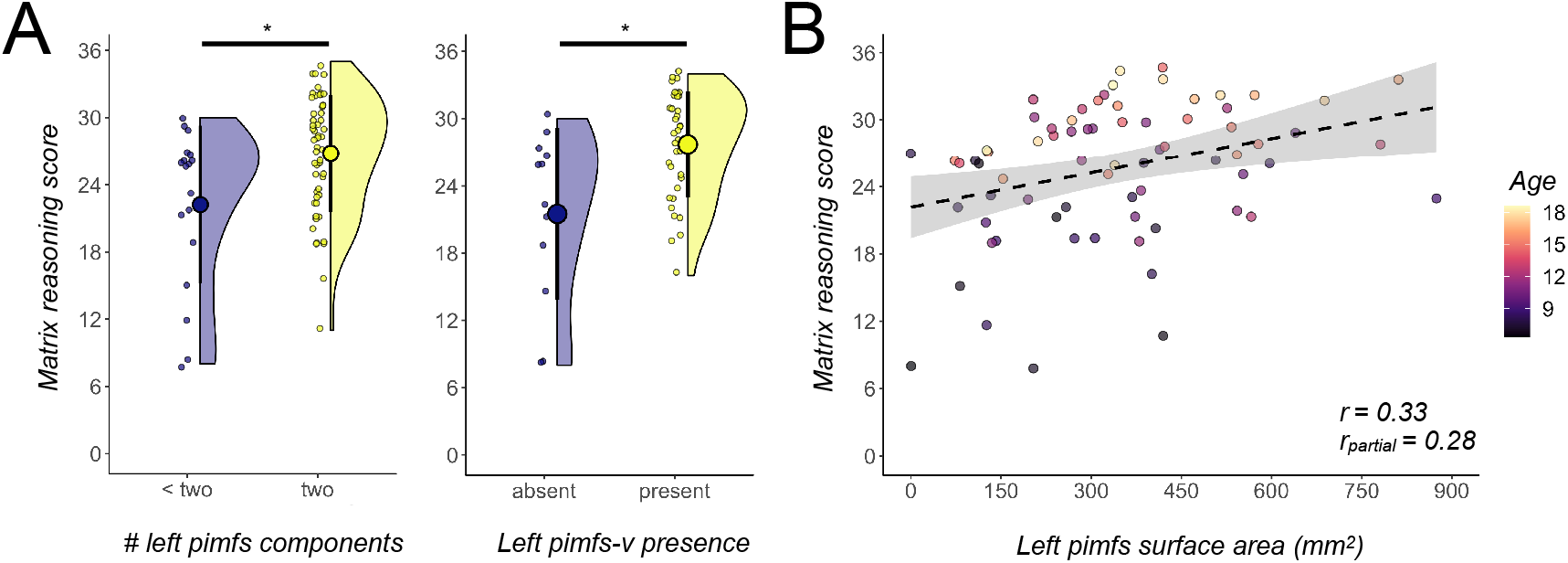
The presence and prominence of the para-intermediate middle frontal sulcus are related to reasoning. (**A**). Raincloud plots depicting reasoning score as a function of (left) the number of pimfs components and (right) the presence of the ventral pimfs component in the left hemisphere only. The large dots and error bars represent the mean±std reasoning score, and the violin plots show the kernel density estimate. The smaller dots indicate individual participants. *Left*: Across the whole sample (N=72), those with two pimfs components (N=54) had better reasoning scores than those with only one component (N=18), controlling for age (\**p* = .045). *Right*: Matching subsamples for age and sample size (total N = 48), participants with the left pimfs-v component had better reasoning performance than those without, also controlling for age (\**p* = .012); this group difference was also observed across the full sample (Supplementary Fig. 2). (**B**) Scatterplot visualizing reasoning scores as a function of left pimfs surface area (raw, in mm^2^), controlling for age. The best fit line, ±95% confidence interval, correlation coefficient (r), and r_partial_ from the regression are included. The smaller dots represent individual participants, colored by age (darker = younger; lighter = older). In the left hemisphere, the surface area of the pimfs is positively associated with reasoning (*p* = .022), even after controlling for age.

Due to differences in the sample size and age distribution of the two groups (median(sd)_present_ = 11.78(3.52), median(sd)_absent_ = 8.81(4.22)), we sought to further confirm the effect of presence/absence of the left pimfs-v on reasoning performance. To this end, we employed variable ratio matching (Materials and Methods) to create an age-matched sample that consisted of the original 12 participants without a left pimfs-v and the 36 age-matched participants with a left pimfs-v (mean age = 10.66, eCDF). A weighted regression in the matched sample with left pimfs-v presence and age as predictors of reasoning revealed that the presence of the left pimfs-v remained significant (ß = 3.69, t = 2.61, *p* = .012; Fig. 2A, right). Critically, this model explained significantly more variance than a model with age alone in the same sample (pimfs: R^2^_adj_ = 0.51, *p* < .001; age: R^2^_adj_ = 0.45, *p* < .001; model comparison: *p* = .012).

Finally, pimfs morphology was also behaviorally relevant when examined with a continuous metric (total surface area of the sulcus), albeit to a lesser degree than the discrete metric of presence/absence of a specific component. Specifically, a linear regression (with age included as a covariate), revealed that the surface area of the left pimfs (ß = 0.01, t = 2.35, *p* = .022) was positively associated with reasoning (Fig. 2B); this relationship was marginal (ß = 0.01, t = 1.85, p = .068) in the right hemisphere. Whereas the discrete models examining sulcal components explained significantly more variance in reasoning than age alone, this model explained marginally more variance than age alone (pimfs: R^2^_adj_ = .49, *p* < .001; age: R^2^_adj_ = .46, *p* < .001; model comparison: *p* = .071). A repeated K-fold (5-fold, 10 repeats) and leave-one-out cross-validation (looCV) confirmed the predictiveness of the pimfs model (*5-fold:* R^2^ = 0.51, RMSE = 4.33; *looCV:* R^2^ = 0.45, RMSE = 4.45), which was only slightly better than age alone (*5-fold:* R^2^ = 0.49, RMSE = 4.40; *looCV:* R^2^ = 0.43, RMSE = 4.50). Further, when normalizing left pimfs surface area by the total surface area of the PFC, the relationship was marginal (*p* = .075). Thus, the presence or absence of pimfs-v may be more directly related to reasoning than pimfs surface area, which is related to PFC surface area more broadly.

## Discussion

Our results reveal that the presence of the left pimfs-v was associated with better reasoning performance. This finding contributes to mounting evidence that the presence or absence of sulci relates to complex cognitive skills (Amiez et al., 2018; Borst et al., 2014; Buda et al., 2011; Cachia et al., 2014; Fornito et al., 2006, 2004; Huster et al., 2011, 2009; Whittle et al., 2009). Crucially, this relationship was not observed for processing speed or working memory, which are theorized to support this high-level cognitive ability (Ferrer et al., 2013; Fry and Hale, 2000). Relatedly, we have recently found that the depth of numerous PFC sulci in the left hemisphere – but not the pimfs – is related to working memory manipulation (Yao et al., in revision). Thus, this brain-behavior relationship does not generalize to another challenging cognitive task. It should also be noted that this result was observed across a large developmental age range (6-18 years old). Future work should seek to determine whether this effect holds longitudinally and into adulthood (Borst et al., 2014; Huster et al., 2011, 2009) – or if this relationship is specific to the time period when higher-level cognitive skills are being acquired.

Relating to previous work, the pimfs-v may be adjacent to rostrolateral PFC, a region that has been heavily implicated in reasoning (e.g., Assem et al., 2020; Christoff et al., 2001; Hartogsveld et al., 2018; Urbanski et al., 2016; Vendetti and Bunge, 2014). However, confirmation of this association awaits individual-level analysis of overlap between pimfs-v and RLPFC neural activation. Considering that the presence of sulci have also been shown to influence the organization of functional networks and task-based activation (Amiez et al., 2013; Amiez and Petrides, 2014; Lopez-Persem et al., 2019), future work should also test whether variations in pimfs incidence are related to functional network organization, as well as task-based activation in LPFC.

The sulcal metrics examined here showed significant effects for left pimfs, with trend-level effects in the right hemisphere. Conversely, we previously showed that sulcal depth of right but not left pimfs (averaged across dorsal and ventral components) was related to reasoning (Voorhies et al., 2021). Both hemispheres have been implicated in reasoning, although there is evidence for functional dissociations between them (Bunge et al., 2009; Goel, 2019; Vendetti et al., 2015). The reason for this hemispheric double-dissociation is not yet clear; perhaps it relates to differential developmental trajectories of, and dynamic relations between, the two hemispheres (Toga and Thompson, 2003), which can be further explored in future research.

Mechanistically, differences in sulcal patterning are hypothesized to be related to the underlying white matter connectivity and broadly speaking, that cortical folding patterns are generally optimized with regards to efficiency of communication between brain regions (Van Essen, 2020, 1997; White et al., 2010; Zilles et al., 2013). Thus, the presence and prominence of the pimfs may result in more efficient neural communication compared to the absence of the pimfs. Additionally, or alternatively, relationships among tertiary sulci, brain function, and behavior could relate to alterations of local cytoarchitecture (Amiez et al., 2021). Individual differences in the presence and prominence of tertiary sulci in association cortices may reflect variability in the rates of growth of adjacent cytoarchitectural regions (e.g., Fernández et al., 2016). Thus, the presence or absence of the pimfs could also reflect differences in local architecture which could, in turn, represent differences in local neural circuits reflecting local computational power supporting reasoning – a multiscale, mechanistic relationship which can be explored in future research.

In conclusion, we have uncovered that the presence of a specific sulcus in LPFC may contribute to the development of reasoning. These findings do not imply a deterministic relationship, for two reasons. First, other neuroanatomical features in LPFC and elsewhere have also been linked to reasoning during development, including sulcal depth (Voorhies et al., 2021), white matter microstructure (Wendelken et al., 2017), and, in some samples, cortical thickness (e.g., Leonard et al., 2019). Thus, a goal of future research is to work toward developing a comprehensive, unifying model that integrates these and any other neuroanatomical features, yet to be identified, that contribute to the development of reasoning. The second reason we do not mean to suggest that the presence or absence of this sulcus determines reasoning ability is that there is evidence of experience-dependent plasticity in reasoning and underlying brain circuitry (e.g., Mackey et al., 2013). Nonetheless, the present findings underscore the behavioral relevance of cortical folding patterns, providing novel insights into one particular LPFC sulcus that exhibits prominent individual differences.

## Materials and Methods

### Participants

The present study consisted of 72 typically developing participants between the ages of 6-18 (mean ± std age = 12.105 ± 3.767 years old, including 30 individuals identified by caregivers as female) that were randomly selected from the Neurodevelopment of Reasoning Ability (NORA) dataset (Ferrer et al., 2013; Wendelken et al., 2017, 2016, 2011). Additional demographic and socioeconomic information is included in Supplementary Table 1. 61 of these participants were also included in prior research on sulcal depth (Voorhies et al., 2021). All participants were screened for neurological impairments, psychiatric illness, history of learning disability, and developmental delay. All participants and their parents gave their informed assent/consent to participate in the study, which was approved by the Committee for the Protection of Human Participants at the University of California, Berkeley.

### Data acquisition

#### Imaging data

fMRI data were collected on a Siemens 3T Trio system at the University of California Berkeley Brain Imaging Center. High-resolution T1-weighted MPRAGE anatomical scans (TR = 2300 ms, TE = 2.98 ms, 1 × 1 × 1 mm voxels) were acquired for cortical morphometric analyses.

#### Behavioral data

All 72 participants completed a matrix reasoning task (WISC-IV), which is a widely used measure of abstract, nonverbal reasoning (Ferrer et al., 2013; Wendelken et al., 2016). Two additional control measures were included when available: processing speed (N = 71) and verbal working memory (N = 64). Reasoning performance was measured as a total raw score from the WISC-IV Matrix reasoning task (Wechsler, 1949; mean±std = 25.65±6.01). Matrix reasoning is an untimed subtest of the WISC-IV in which participants are shown colored matrices with one missing quadrant. The participant is asked to “complete” the matrix by selecting the appropriate quadrant from an array of options. Previous factor analysis in this dataset (Ferrer et al., 2013) showed that the Matrix reasoning score loaded strongly onto a reasoning factor that included three other standard reasoning assessments consisting of the *Block Design* subtest of the Wechsler Abbreviated Scale of Intelligence (WASI; Wechsler, 1999), as well as the *Analysis Synthesis* and *Concept Formation* subtests of the Woodcock-Johnson Tests of Achievement (Woodcock et al., 2001).

Processing speed was computed from raw scores on the Cross Out task from the Woodcock-Johnson Psychoeducational Battery-Revised (WJ-R; Brown et al., 2012). In this task, the participant is presented with a geometric figure on the left followed by 19 similar figures. The participant places a line through each figure that is identical to the figure on the left of the row. Performance is indexed by the number of rows (out of 30 total rows) completed in 3 minutes (mean±std = 22.1±6.75). Cross Out scores are frequently used to estimate processing speed in developmental populations (Kail and Ferrer, 2007; McBride-Chang and Kail, 2002).

Verbal working memory was measured via raw scores of the Digit Span task from the 4th edition of the Wechsler Intelligence Scale for Children (WISC-IV; (Wechsler, 1949). The Digits Forward condition of the Digit Span task taxes working memory maintenance, whereas the Backward condition taxes both working memory maintenance and manipulation. In Digits Forward, the experimenter reads aloud a sequence of single-digit numbers, and the participant is asked to immediately repeat the numbers in the same order; in Digits Backward, they are asked to immediately repeat the numbers in the reverse order. The length of the string of numbers increases after every two trials. The Forwards task has eight levels, progressing from 2 to 9 digits. The Backwards task has seven levels, from 2 to 8 digits. Participants are given a score of 1 for a correct answer or a 0 for an incorrect answer. Testing on a given task continues until a participant responds incorrectly to both trials at a given level, after which the experimenter recorded a score out of 16 for Digits Forward (16 total trials; mean±std = 9.03±2.24) and a score out of 14 for Digits Backward (14 total trials; mean±std = 5.84±2.12).

### Morphological analyses

#### Cortical surface reconstruction

All T1-weighted images were visually inspected for scanner artifacts. FreeSurfer’s automated segmentation tools (Dale et al., 1999; Fischl and Dale, 2000; FreeSurfer 6.0.0) were used to generate cortical surface reconstructions. Each anatomical T1-weighted image was segmented to separate gray from white matter, and the resulting boundary was used to reconstruct the cortical surface for each participant (Dale et al., 1999; Wandell et al., 2000). Each reconstruction was visually inspected for segmentation errors, and these were manually corrected when necessary.

Cortical surface reconstructions facilitate the identification of shallow tertiary sulci compared to post-mortem tissue for two main reasons. First, T1 MRI protocols are not ideal for imaging vasculature; thus, the vessels that typically obscure the tertiary sulcal patterning in post-mortem brains are not imaged on standard resolution T1 MRI scans. Indeed, indentations produced by these smaller vessels that obscure the tertiary sulcal patterning are visible in freely available datasets acquired at high field (7T) and micron resolution (100–250 μm; Edlow et al., 2019; Lüsebrink et al., 2017). Thus, the present resolution of our T1s (1 mm isotropic) is sufficient to detect the shallow indentations of tertiary sulci but is not confounded by smaller indentations produced by vasculature. Second, cortical surface reconstructions are made from the boundary between gray and white matter; unlike the outer surface, this inner surface is not obstructed by blood vessels (Weiner, 2019; Weiner et al., 2018).

#### Defining the presence and prominence of the para-intermediate middle frontal sulcus

We first manually defined the pimfs within each individual hemisphere in *tksurfer* (Miller et al., 2021b). Manual lines were drawn on the *inflated* cortical surface to define sulci based on the most recent schematics of pimfs and sulcal patterning in LPFC by Petrides (2019), as well as by the *pial* and *smoothwm* surfaces of each individual (Miller et al., 2021b). In some cases, the precise start or end point of a sulcus can be difficult to determine on a surface (Borne et al., 2020). Thus, using the *inflated, pial*, and *smoothwm* surfaces of each individual to inform our labeling allowed us to form a consensus across surfaces and clearly determine each sulcal boundary. For each hemisphere, the location of the pimfs was confirmed by three trained independent raters (E.H.W., W.I.V., J.K.Y.) and finalized by a neuroanatomist (K.S.W.). Although this project focused on a single sulcus, it took the manual identification of all LPFC sulci (2448 sulcal definitions across all 72 participants) to ensure the most accurate definitions of the pimfs components (for descriptions of these LPFC sulci see (Miller et al., 2021a, 2021b; Petrides, 2019; Voorhies et al., 2021; Yao et al., in revision).

Individuals typically have three to five tertiary sulci within the middle frontal gyrus (MFG) of the lateral prefrontal cortex (Miller et al., 2021a, 2021b; Voorhies et al., 2021; Yao et al., in revision). The posterior MFG contains three of these sulci, which are present in all participants: the anterior (pmfs-a), intermediate (pmfs-i), and posterior (pmfs-p) components of the posterior middle frontal sulcus (pmfs; (Miller et al., 2021a, 2021b; Voorhies et al., 2021; Yao et al., in revision). In contrast, the tertiary sulcus within the anterior MFG, the para-intermediate middle frontal sulcus (pimfs), is variably present. A given hemisphere can have zero, one, or two pimfs components (Voorhies et al., 2021; Yao et al., in revision).

Drawing from criteria outlined by Petrides (2019, 2013), the dorsal and ventral components of the para-intermediate middle frontal sulcus (pimfs-d and pimfs-v) were generally defined using the following two-fold criterion: i) the sulci ventrolateral to the horizontal and ventral components of the intermediate middle frontal sulcus, respectively, and ii) superior and/or anterior to the mid-anterior portion of the inferior frontal sulcus. The location of each indentation was cross-checked using the *inflated, pial*, and *smoothwm* surfaces. We first confirmed the accuracy of this criterion by applying it to the individual participants with two identifiable pimfs sulci. Next, we extended this criterion to label the cases in which an individual only had one component. We then compared incidence rates between components and hemispheres with a chi-squared and Fischer exact test, respectively.

We quantified the prominence of the pimfs as its surface area (in mm^2^). The surface area values for each pimfs label were extracted using the *mris_anatomical_stats* function that is included in FreeSurfer (Fischl and Dale, 2000). For those with two pimfs components, the surface area was extracted as a sum of both components together (via a merged label with *mris_mergelabel* function (Dale et al., 1999) and for each individual component separately. We also considered normed values. To normalize pimfs surface area by the surface area of the PFC, we automatically defined the PFC in both hemispheres of each participant with the *mris_annot2label --lobesStrict* function and then extracted surface area values with the *mris_anatomical_stats* function (Fischl and Dale, 2000).

### Behavioral Analyses

#### Relating the presence of the pimfs to reasoning performance

To assess whether the presence of the pimfs in each hemisphere is related to reasoning performance, we first conducted an analysis of covariance (ANCOVA) with number of components in the left and right hemispheres (*two or less than two*) as factors, and assessment age as a covariate. Gender was not considered in these analyses, as it was not associated with either matrix reasoning (*p* = 0.65) or number of components (*left*: *p* = 0.27, *right*: *p* = 0.8). Next, to determine if the presence of a specific pimfs component was related to reasoning performance, we ran a second ANCOVA with left and right hemisphere presence of the pimfs-v and pimfs-d (*yes, no*) as factors and age as a covariate. Although the presence of a left pimfs-d was related to age (*p* = .02), this moderate collinearity did not affect the model results (vif < 5).

#### Control behavioral analyses

To ascertain whether the relationship between left pimfs-v presence and cognition is specific to reasoning performance, or generalizable to other general measures of cognitive processing (Kail and Ferrer, 2007), we tested this sulcal-behavior relationship with two other widely used measures of cognitive functioning: processing speed and working memory maintenance and manipulation. Specifically, we ran three ANCOVAs with left pimfs-v presence (*yes, no*) as a factor and assessment age as a covariate.

#### Matching analysis

Variable ratio matching (ratio = 3:1, min = 1, max = 5) was conducted with the *MatchIt* package in R (https://cran.r-project.org/web/packages/MatchIt/MatchIt.pdf). The optimal ratio parameter was determined based on the calculation provided by (Ming and Rosenbaum, 2000). To accommodate variable-ratio matching, the distance between each member of each group was computed by a logit function:

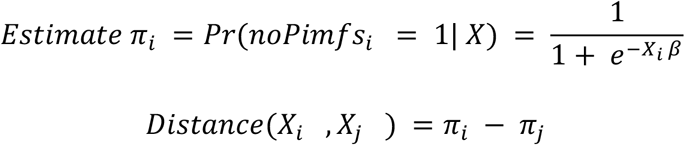

where X is participant age in groups without (*i*) and with (*j*) a pimfs_lh_ventral_. Matches were determined by greedy nearest-neighbor interpolation such that each participant in the smaller group received at least one, and up to five, unique matches from the larger group.

#### Relating the size of the pimfs to reasoning performance

To test if the prominence (surface area) of the pimfs was related to reasoning performance, we implemented a multiple linear regression with surface area of pimfs (combined if two were present) in left and right hemispheres as predictors, while controlling for assessment age. Gender was not included, as it was not related to surface area in either hemisphere (*left*: *p* = 0.16, *right*: *p* = 0.78) and, as noted previously, it was also not related to reasoning performance. Despite there being a significant correlation between age and left pimfs SA (r = 0.24, *p* = .042) and a trending correlation between age and right pimfs SA (r = .20, *p* = .09), this collinearity did not affect the model results (vifs < 4); thus, all predictors were still included. This model was further cross-validated with leave-one-out (looCV) and repeated K-fold (5-fold, 10 repeats) cross-validation methods. Finally, to assess whether prefrontal surface area affected the model, we also ran an exploratory linear regression with normed surface area of the pimfs (by hemispheric PFC surface area) in left and right hemispheres with the covariate assessment age as predictors

#### Model comparison

All nested pimfs models were compared to age alone with Chi-Squared tests.

#### Statistical tests

All statistical tests were implemented in R v4.1.2 (https://www.r-project.org/). Incidence chi-squared tests were carried out with the *chisq.test* function from the R *stats* package. Fisher’s exact tests were carried out with the *fisher.test* function from the R *stats* package. All ANCOVAs were implemented using the *lm* and *Anova* functions from the R *stats* and *cars* packages. Effect sizes for the ANCOVA effects are reported with the *generalized* eta-squared (η2G) metric. Linear models were run using the *lm* function from the R *stats* package. Leave-one-out and K-fold cross-validation analyses were carried out with the *train.control* and *train* functions from the R *caret* package. The effect of each pimfs model was compared to the effect of age alone with the *anova* function from the R *stats* package.

#### Data availability

Data used for this project have been made freely available on GitHub (https://github.com/cnl-berkeley/stable_projects/tree/main/PresAbs_Reasoning). Requests for further information or raw data should be directed to the Corresponding Author, Kevin Weiner.

## Acknowledgments

This research was supported by NICHD R21HD100858 (Weiner, Bunge), NSF CAREER Award 2042251 (Weiner), and an NSF-GRFP fellowship (Voorhies). Funding for original data collection and curation was provided by NINDS R01 NS057156 (Bunge, Ferrer) and NSF BCS1558585 (Bunge, Wendelken). We thank former members of the Bunge laboratory for assistance with data collection, and the families who participated in the original study.

## Author contributions

E.H.W., W.I.V., K.S.W, and S.A.B. designed research; E.H.W., W.I.V., J.K.Y., and K.S.W. performed manual sulcal labeling; E.H.W. and W.I.V. analyzed data; E.H.W., W.I.V., K.S.W., and S.A.B. wrote the paper; all authors edited the paper and gave final approval before submission.

