## Supplementary Materials for "Presence or absence of a prefrontal sulcus is linked to reasoning performance during child development"

**This PDF file includes:**

Supplementary Figs. 1-2
Supplementary Table 1

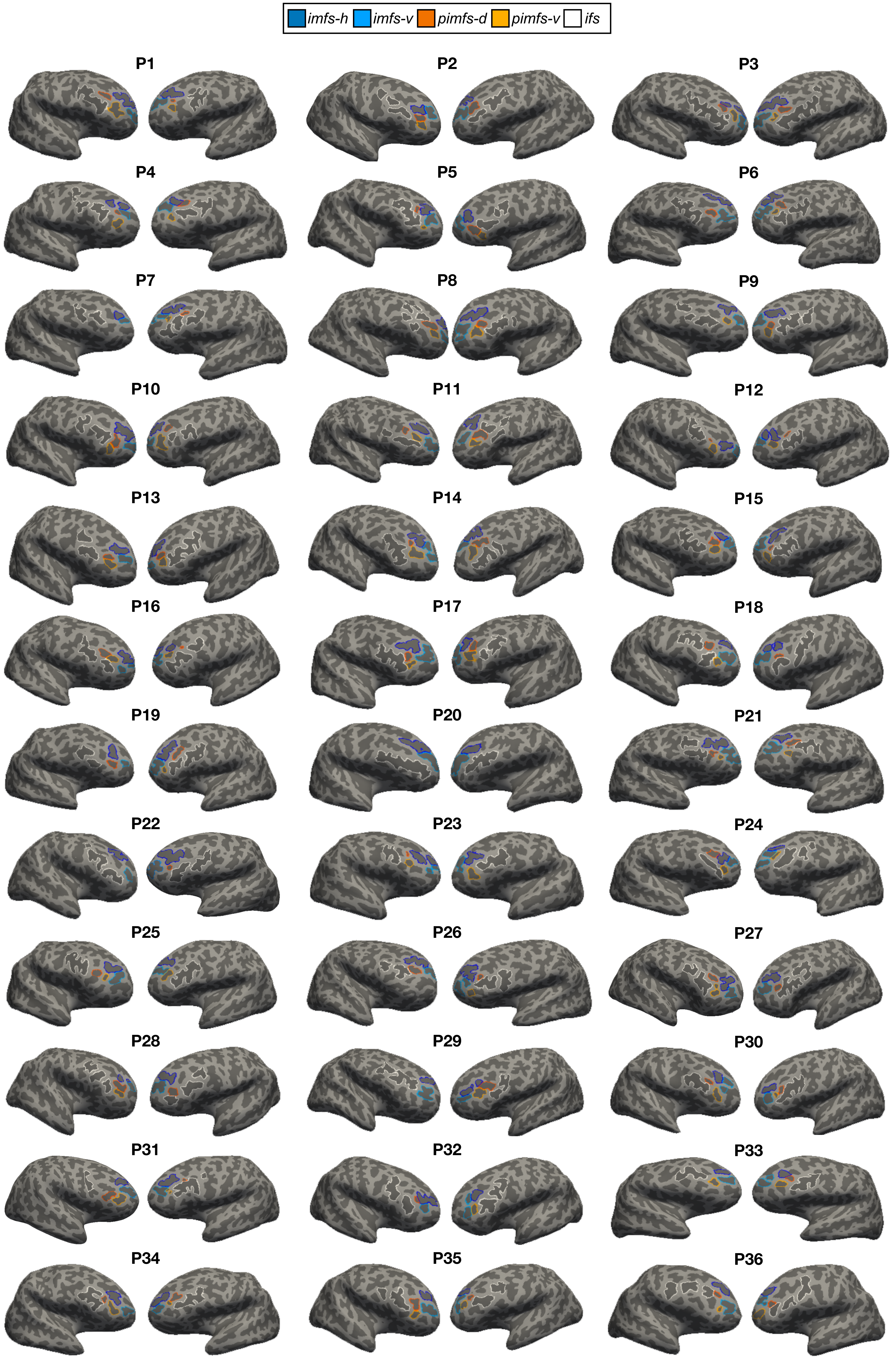
imfs-h
imfs-v
pimfs-d
pimfs-v
ifs
P1
P2
P3
P4
P5
P6
P7
P8
P9
P10
P11
P12
P13
P14
P15
P16
P17
P18
P19
P20
P21
P22
P23
P24
P25
P26
P27
P28
P29
P30
P31
P32
P33
P34
P35
P36

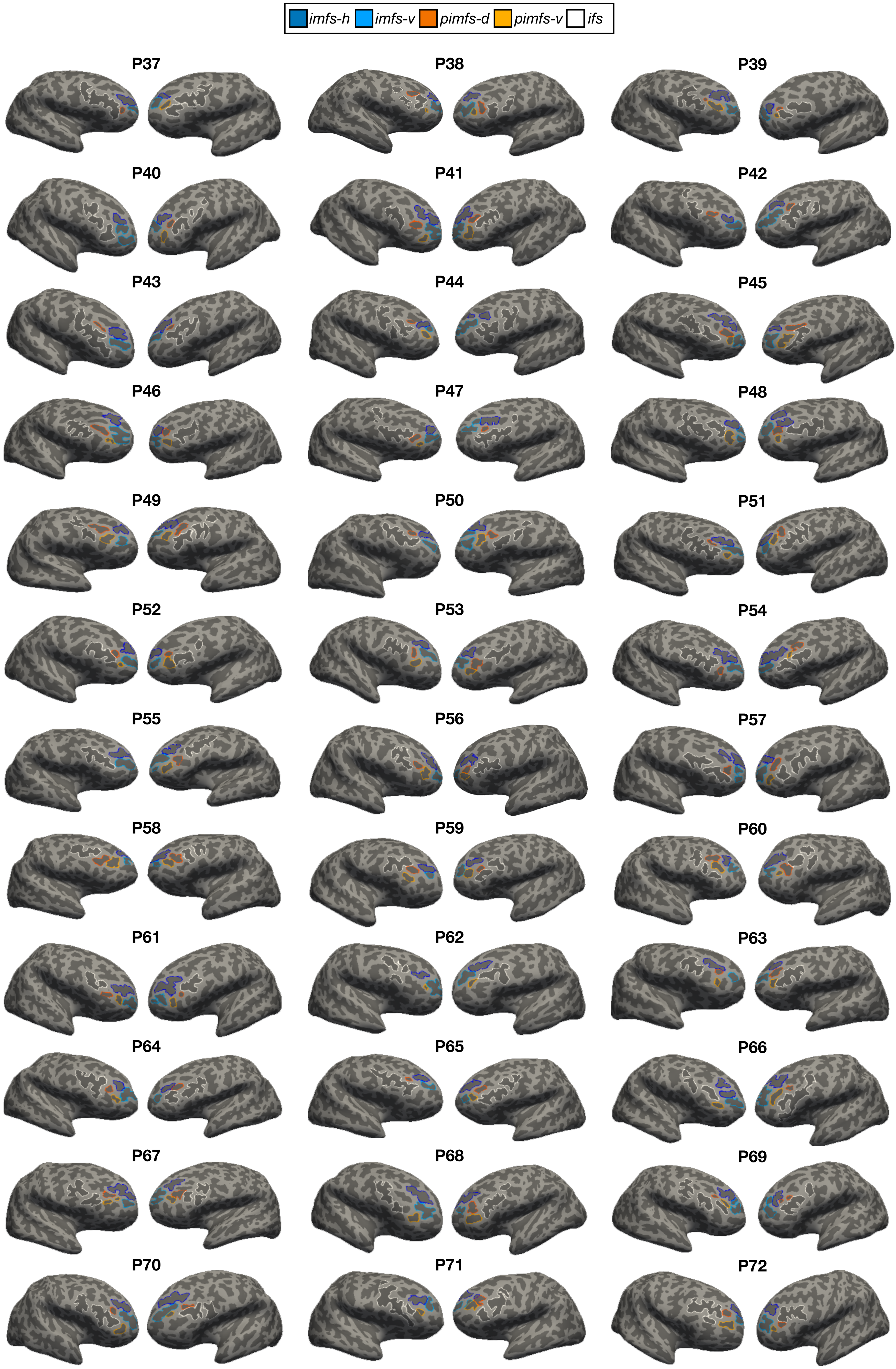
imfs-h
imfs-v
pimfs-d
pimfs-v
ifs
P37
P38
P39
P40
P41
P42
P43
P44
P45
P46
P47
P48
P49
P50
P51
P52
P53
P54
P55
P56
P57
P58
P59
P60
P61
P62
P63
P64
P65
P66
P67
P68
P69
P70
P71
P72

**Supplementary Fig. 1. Manual sulcal labels in the left and right hemispheres of each participant (N=72).** As in Figure 1A, each sulcus is displayed on the inflated cortical surface (surfaces are not to scale) in FreeSurfer 6.0.0 and is colored according to the key at the top. Sulci were defined according to the most recent atlas and criteria by Petrides (2013, 2019; Materials and Methods). All hemispheres have the horizontal (imfs-h; dark blue) and ventral (imfs-v; light blue) intermediate middle frontal sulci and inferior frontal sulcus (ifs; white). The pimfs (orange) is more variable: participants can have zero, one (dorsal (dark) or ventral (light)), or two components (dorsal and ventral).

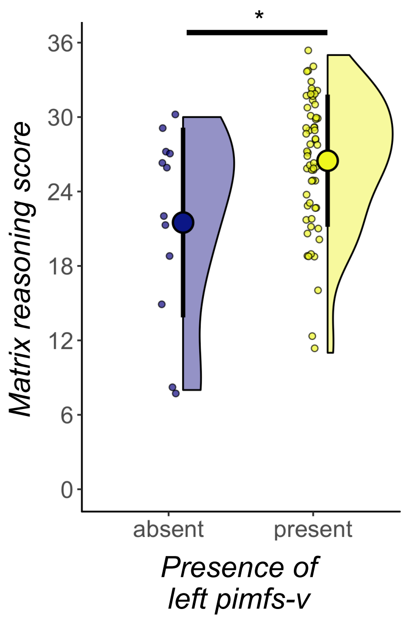


**Supplementary Fig. 2. The presence of the ventral para-intermediate middle frontal sulcus is related to reasoning (whole sample).** Raincloud plot depicting reasoning score as a function of the presence of the ventral pimfs component in the left hemisphere using the whole sample (N=72). The large dots and error bars represent the mean±std reasoning score and the violin shows the kernel density estimate. The smaller dots indicate individual participants. These features are colored (blue and yellow) to distinguish between the two groups. After controlling for age, those with the ventral component in the left hemisphere (*p = .027) had better reasoning scores than those without.

| | Count |
|---|---|
| Racial categories | |
| American Indian/Alaskan Native | 0 |
| Asian/Native Hawaiian/Other Pacific Islander | 5 |
| Black or African American | 4 |
| White | 47 |
| More Than One Race | 14 |
| Unknown or Not Reported | 2 |
| Ethnic categories | |
| Hispanic or Latino | 11 |
| Not Hispanic or Latino | 59 |
| Unknown or Not Reported | 2 |
| Highest degree earned by parent/guardian | |
| High School/GED | 8 |
| Vocational | 1 |
| Associate degree | 6 |
| Bachelor's degree | 18 |
| Master's degree | 15 |
| Doctorate | 4 |
| Professional | 3 |
| Other | 3 |
| None of the above (less than high school) | 1 |
| Unknown or Not Reported | 13 |
| Total household income | |
| $16,000-$24,999 | 2 |
| $25,000-$34,999 | 3 |
| $50,000-$74,999 | 6 |
| $75,000-$99,999 | 8 |
| $100,000-$199,999 | 27 |
| Over $200,000 | 5 |
| Unknown or Not Reported | 21 |

**Supplementary Table. 1. Demographic and socioeconomic information of the child/adolescent sample (N = 72).** All information is parent/guardian reported.
